# Bayesian mixture analysis for metagenomic community profiling

**DOI:** 10.1101/007476

**Authors:** Sofia Morfopoulou, Vincent Plagnol

## Abstract

Deep sequencing of clinical samples is now an established tool for the detection of infectious pathogens, with direct medical applications. The large amount of data generated provides an opportunity to detect species even at very low levels, provided that computational tools can effectively interpret potentially complex metagenomic mixtures. Data interpretation is complicated by the fact that short sequencing reads can match multiple organisms and by the lack of completeness of existing databases, in particular for viral pathogens. This interpretation problem can be formulated statistically as a mixture model, where the species of origin of each read is missing, but the complete knowledge of all species present in the mixture helps with the individual reads assignment. Several analytical tools have been proposed to approximately solve this computational problem. Here, we show that the use of parallel Monte Carlo Markov chains (MCMC) for the exploration of the species space enables the identification of the set of species most likely to contribute to the mixture. The added accuracy comes at a cost of increased computation time. Our approach is useful for solving complex mixtures involving several related species. We designed our method specifically for the analysis of deep transcriptome sequencing datasets and with a particular focus on viral pathogen detection, but the principles are applicable more generally to all types of metagenomics mixtures. The work is implemented as a user friendly R package, available from CRAN: http://cran.r-project.org/web/packages/metaMix.

## Introduction

Metagenomics can be defined as the analysis of a collection of DNA or RNA sequences originating from a single sample. In practice, its scope is broad and includes the analysis of a diverse set of samples such as gut microbiome (Qin et al., 2010), (Minot et al., 2011), environmental (Mizuno et al., 2013) or clinical (Willner et al., 2009), (Negredo et al., 2011), (McMullan et al., 2012) samples. Among these applications, the discovery of viral pathogens is clearly relevant for clinical practice (Fancello et al., 2012), (Chiu, 2013). The traditional process of characterizing a virus through potentially difficult and time consuming culture techniques is being revolutionized by advances in high throughput sequencing. Potential benefits of sequence driven methodologies include a more rapid turnaround time (Quail et al., 2012), combined with a largely unbiased approach in species detection, including the opportunity for unexpected discoveries.

The analysis of shotgun sequencing data from metagenomic mixtures raises complex computational challenges. Part of the difficulty stems from the read length limitation of existing deep DNA sequencing technologies, an issue compounded by the extensive level of homology across viral and bacterial species. Another complication is the divergence of the microbial DNA sequences from the publicly available references. As a consequence, the assignment of a sequencing read to a database organism is often unclear. Lastly, the number of reads originating from a disease causing pathogen can be low (Barzon et al., 2013). The pathogen contribution to the mixture depends on the biological context, the timing of sample extraction and the type of pathogen considered. Therefore, highly sensitive computational approaches are required.

A first analytical problem is read classification, that is the assignment of a given sequencing read to a species. Several tools have been developed and these belong to two broadly defined classes: composition-based and similarity-based approaches. The read classification based on sequence composition relies on the intrinsic features of the reads, such as CG content or oligonucleotide distributions. Methods include PhyloPythia (McHardy et al., 2007), Phymm (Brady and Salzberg, 2009), MetaCluster (Yang et al., 2010). These tend to focus on major classes in a dataset and may not perform well on low-abundance populations (Kunin et al., 2008). Additionally, results are usually reliable for longer reads only (Dröge and McHardy, 2012).

Similarity based methods, using homology search algorithms such as BLAST (Altschul et al., 1990), are considered the most sensitive methods for read classification (Brady and Salzberg, 2009). One of the most popular tools using the output of a similarity search algorithm is MEGAN (Huson et al., 2007).

MEGAN addresses ambiguous matches by assigning reads that have multiple possible assignments to several species, to the taxonomic group containing all these species, or else their lowest common ancestor (LCA). This approach is accurate on a higher taxonomic level. However, it is lacking a formal solution to resolving ambiguous matches. A weakness of the similarity based methods is that a long tail of species, each supported only by a few reads can appear in the results. This results from the classification being decided one read at a time, in contrast to considering all reads simultaneously. Hybrid methods combining composition and similarity information such as PhymmBL (Brady and Salzberg, 2009) and RITA (MacDonald et al., 2012) also tackle one read at a time.

Methods focused on the statistical inference of the set of present species as well as the estimation of their relative proportions, incorporate knowledge from all reads to assign each individual read to a species. From a statistical standpoint, this identification and quantification question can be thought of as an application of mixture models. These ideas have been applied in the metagenomics context in frequentist (GRAMMy (Xia et al., 2011)) and Bayesian (Pathoscope (Francis et al., 2013)) settings. GRAMMy formulates the problem as a finite mixture model, using the Expectation-Maximization (EM) algorithm to estimate the relative genome abundances. Pathoscope refines this process by penalizing reads with ambiguous matches in the presence of reads with unique matches and enforcing parsimony within a Bayesian context. Both methods work with unassembled sequence data and they are not currently setup to incorporate an initial short read assembly step, which could be achieved by assigning a higher weight to contigs formed by multiple reads.

Fitting a mixture model is useful for the species relative abundance estimation, as well as the read to species assignment. A related but distinct question concerns the set of species which should be included in the mixture model. This question is closely related to the biological question of asking what species are present in the mixture. Including all species flagged as potential matches by the read classification can introduce a large number of species, often in the low thousands. Mixture models will, in this situation, identify a large number of species at low levels. This interpretation is appropriate in some applications. In many other cases, the expectation is that the underlying species set should be parsimonious and that some divergence with database species or sequencing errors can explain a large fraction of the non matching reads.

Hence, a better statistical formulation of the community profiling problem is the exploration of the candidate organisms state-space. In this context, non nested models can be compared based on their marginal likelihood. Within this Bayesian framework, readily interpretable probabilities, such as the posterior probabilities of species sets can be used to quantify the support for a species in the mixture. Finally, more complex hypotheses testing for example the number of viral species or the joint presence of two distinct organisms can be investigated.

The main challenge behind such a formulation is computational. Even with a relatively small number of species to consider, the number of subsets of this space that could explain the mixture grows exponentially. Efficient computational strategies are required to make this problem tractable. Here we show that this inference can be achieved for modern scale metagenomics datasets. Our strategy is based on parallel tempering, a Monte Carlo Markov Chain technique, using parallel computing to speed up the inference. We implemented these ideas in a user friendly R package called metaMix. metaMix produces posterior probabilities for various models as well as the relative abundances under each model. We demonstrate its potential using datasets from clinical samples as well as benchmark metagenomic datasets.

## Results

We first applied metaMix on a popular benchmark dataset, for which the community composition and the read assignment is known. We then analyzed RNA-Seq datasets from two clinical samples that were generated for diagnostic purposes. We compare our results with the ones produced by MEGAN version 5.3 and Pathoscope 2.0. Both methods are similarity-based. This property and more specifically their flexibility to work with BLASTx output, makes them better candidates for viral discovery compared to composition-based methods. From the mixture model methods, we have chosen Pathoscope. We were also interested in comparing our results to the ones by GRAMMy, which was the first similarity-based method to use the idea of the mixture model. However, GRAMMy is designed for nucleotide-nucleotide comparisons (BLASTn), which is suboptimal for viral discovery. GRAMMy also only considers unassembled reads and requires that these are of the same fixed length. For these reasons, GRAMMy was not included in the comparison. Default parameters were used for all methods, unless stated otherwise.

For the metaMix output, we reported organisms with a posterior probability greater than 0.9. The metaMix read support parameter r, which essentially sets the sensitivity/specificity of the method, has an impact on the number of reported species. A large r value can result in the method merging together strains that are differentiated by fewer reads than r. On the other hand a low r can have the opposite effect, whereby the methods splits a strain into two or more strains, by moving a few reads from one strain to a very similar one with which they have equally good matches.

The user’s choice for this key parameter r should be informed by the biological context. As an example, for the typical human clinical sample where the sample collection might have occurred some time after the infection has taken place, a low value in order to adopt a sensitive approach is reasonable. Hence, for viral identification in human clinical samples, a low and sensitive value (r = 10) is a reasonable choice. In a highly complex environmental metagenomic community where there is a plethora of species of similar abundances, the choice becomes less straightforward especially in the case of closely related strains. We set the default value for general community profiling in environmental samples to be r = 30. We also compare the output of metaMix for different values of this parameter. Supplementary Text 1 presents a detailed discussion on these settings as well as practical considerations.

## FAMeS simHC Dataset – Closely Related Strains

The FAMeS artificial datasets (http://fames.jgi-psf.org/description.html), are simulated metagenomic datasets composed of random reads from 113 isolate microbial genomes present in IMG (Integrated Microbial Genomes) and sequenced at the DOE Joint Genome Institute. They are a popular choice to use as benchmark datasets for various metagenomics methods. Their suitability stems from the fact that the number of species that form the metagenomic community is known as well as their relative abundances. The FAMeS datasets have been designed to model real metagenomic communities in terms of complexity and phylogenetic composition.

There are three datasets: simHC, simMC, simLC corresponding to high, medium and low complexity of the metagenomic community respectively. The three methods were applied to simHC, as this is the highest complexity dataset, with many closely related strains with similar abundances and no dominant species. The lowest abundance is 255 reads out of 118,000 reads. The bioinformatics processing in this instance consisted of a BLASTn comparison to all NCBI bacterial genomes (ftp://ftp.ncbi.nlm.nih.gov/genomes/Bacteria/all.fna.tar.gz). The number of genomes mapped, retrieved from the the BLASTn output was ~2,500.

As discussed below, metaMix outperforms Pathoscope and MEGAN in the community profiling task (Table 1) and consequently in the relative abundance estimation (Table 2).

**Table 1.** Number of species identified for the FAMeS simHC dataset.

| True | MEGAN | Pathoscope | metaMix |
| --- | --- | --- | --- |
| 113 | 232 | 165 | 116 |

**Table 2.** Measures of estimation accuracy of relative abundances for the FAMeS simHC dataset.

| Method | RRMSE | AVGRE |
| --- | --- | --- |
| metaMix | 17 | 8.5 |
| Pathoscope | 36.6 | 29.7 |
| MEGAN | 35.9 | 18 |

### metaMix

To limit the complexity of the fit, we used the two step procedure described in the Methods and fully implemented in metaMix. We first fitted the mixture model with the complete set of 2,500 species and a limited run length of 500 iterations. Based on this analysis, we identified 1,312 species supported by at least one read and explored this state space. To limit the computational time, we also considered a stronger approximation, including only the 374 potential species supported by at least 10 sequencing reads. Both approaches generated similar results, albeit the more complex one with 1,312 potential species required the quadruple of the computation time (12h instead of 3h). metaMix identified 116 species, detecting successfully all the members of the metagenomic community (Supplementary Table S1). These were detected on the strain level except in four instances where a different strain of the same species, or different species within the same genus was detected. Four species were identified and not in the simulated dataset, hence can be considered as false positives (Supplementary Table S1).

### Pathoscope

Pathoscope identified 47 species. Of these 42 are members of the metagenomic community. 42 are the exact same strain, while 3 are either the same species but different strain or same genus but different species. However it fails to detect 68 species that are actually present in the mixture. Tuning the parameter that enforces the parsimonious results (any thetaPrior greater than 10), thereby removing the unique read penalty, Pathoscope behaves as a standard mixture model and identifies 165 species. With these settings, it identifies all but one members of the community. The organisms are identified at the strain level, except in three instances where it identified different species within the same genus. The major interpretation issue is the presence of a long tail of species (54 species) that are actually not present in the mixture (Supplementary Table S1). Pathoscope produced the results in one minute.

### MEGAN

MEGAN identified 232 species. It discovered all original species of the community on the strain level, except for 9 instances where it identified the lowest common ancestor (LCA). Aside from the lack of strain or species specificity for 8% of the community members, the main issue is the long tail of false positives species. In the species summary provided by MEGAN, there are 119 species which are not actually present, but supported by a sufficient number of reads (default value: 50 reads) for MEGAN to include these in the output. It finished the computations in less than one minute.

### Relative Abundances

The primary aim for metaMix is to be a diagnostic tool and to answer whether a species is present or absent from the mixture we study. As a secondary aim, we are also interested in estimating accurately the relative abundance of the present organisms. We can assess the abundance estimates produced by the methods by using error measures such as the relative root mean square error, RRMSE and the average relative error, AVGRE. For all methods, when the exact strain was not identified but the correct species or genus was, we used this abundance. 
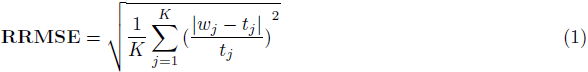

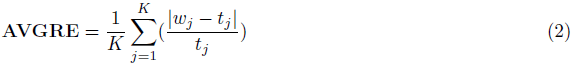
where *t_j_*. is the true abundance of species *j* and *w_j_* the estimated abundance. metaMix produces the most accurate abundance estimates and the results are summarized in Table 2.

### Importance of Read Support Parameter

We then assessed the importance of the read support parameter r on the output of metaMix. We ran metaMix on the benchmark simHC FAMeS dataset with r = {10, 20, 30, 50} reads (Table 3, Figure 1). We observe that as r decreases, a few more related strains from the reference database that are not in the community are retained in the output. As r increases two similar strains are merged into one.

**Figure 1:**
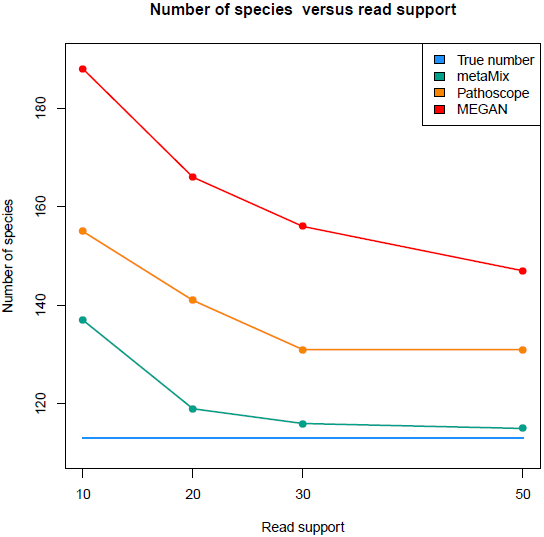
Number of species in the simulated simHC FAMeS dataset detected by metaMix, Pathoscope and MEGAN, as a function of the minimum number of reads required for each species to appear in the output. For metaMix that is r={10, 20, 30, 50} reads, for Pathoscope thetaPrior> 7+ post-run threshold ={10, 20, 30, 50} reads, for MEGAN ”Min Support” + post-run threshold ={10, 20, 30, 50} reads.

**Table 3.**
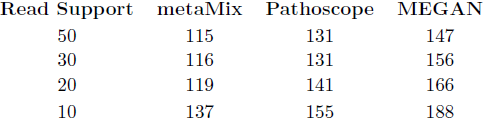
Number of species in the simHC FAMeS dataset by metaMix, Pathoscope and MEGAN

We compared these results with the output of Pathoscope and MEGAN. None of these methods have a read support parameter serving the same purpose as in metaMix, so we tuned the most relevant parameters in these tools. Pathoscope has a thetaPrior parameter that enforces a unique read penalty. This parameter represents the read pseudocounts for the non-unique matches and the default setting is zero which allows for non informative priors. Using the default setting Pathoscope identifies 47 taxa. When thetaP’s value is in (1,7) it identifies 22 taxa, while with thetaP> 7 it identifies 165. With this latter setting which is the one we chose for the comparison, Pathoscope behaves as a standard mixture model.

MEGAN has a ”Min Support” parameter which sets a threshold for the number of reads that must be assigned to a taxon so that it appears in the result. Any read assigned to a taxon not having the required support is pushed up the taxonomy until a taxon is found that has sufficient support. We used Min support = {10, 20, 30, 50} reads. The respective number of taxa in the summary files were 250, 243, 236, 232.

We then also applied a post-run read count threshold to both methods’ output summary. We set the threshold for 10,20,30,50 reads respectively, disregarding taxa that have less than that number of reads assigned to them. In all instances metaMix produces a community profile closer to the real one (Table 3, Figure 1) Pathoscope finds ~ 20 more false positives while MEGAN ~ 40 more compared to metaMix at the same read support level.

The simHC from the FAMeS datasets is complex and distinct from typical human clinical samples, putting aside gut microbiome analysis. The differences are the large number of organisms, the presence of closely related strains of similar abundances, as well as the lack of viruses. Nevertheless, it is an essential dataset to use as a benchmark for examining the performance of the methods in a situation of closely related strains in the sample.

## Human Clinical Sample – Low Viral Load

### Protein Reference Database

For the analysis of human clinical samples, we use a custom reference database that combineds viral, bacterial, human and mouse RefSeq proteins. All viruses are used (ftp://ftp.ncbi.nlm.nih.gov/refseq/release/viral/viral.1.protein.faa.gz) as well as all the bacteria of the human microbiome, according to ftp://ftp.ncbi.nih.gov/genomes/HUMAN_MICROBIOM/Bacteria/all.faa.tar.gz.

To test metaMix in a clinical setting with a low viral load, we used a brain biopsy RNA-Seq dataset from an undiagnosed encephalitis patient (UCL Hospital, data provided as part of a collaboration with Professor Breuer, UCL). Total RNA was purified from the biopsy and polyA RNA was separated for sequencing library preparation. The Illumina MiSeq instrument generated 20 million paired-end reads. We processed the raw data using the bioinformatics pipeline described in Methods. The processed dataset consisted of ~ 75, 000 non-host reads and contigs. Based on the BLASTx output there were 1,298 potential species.

### metaMix

Following the initial processing, we used metaMix for species identification and abundance estimation. The resulting species profile is shown in Table 4; the 13 metaMix entries correspond to 10 species. The most abundant organism was the ϕX174 bacteriophage, which is routinely used for deep-sequencing quality control. More interestingly, we identified an astrovirus. Five short assembled contigs (44 reads) with length ranging between 167bp and 471bp and two non-assembled reads were assigned to the *Astrovirus VA1* with a probability score of 1 (Figure 2). metaMix also identified a number of bacteria supported by a few reads. These are either known laboratory reagent contaminants or human skin associated contaminants (Salter et al., 2014). The analysis completed in 29 minutes.

**Figure 2:**
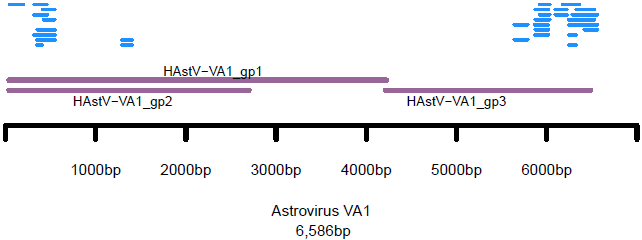
Human clinical sample – novel virus. The reads (blue lines) assigned by metaMix to Astrovirus VA1, aligned to its genome. The purple lines represent the genes of the virus.

**Table 4.** Human clinical sample – novel virus. Comparison of community profile: metaMix – Pathoscope.

| Taxon Identifier | Scientific Name | metaMix |  | Pathoscope |
| --- | --- | --- | --- | --- |
|  |  | Assigned Reads | Posterior Probability | Final best hit read numbers |
| 374840 | *Enterobacteria phage phiX174 sensu lato* | 60449 | 1 | 65327 |
| NA | *unknown* | 10254 | 1 | NA |
| 9606 | *Homo sapiens* | 216 | 1 | 554 |
| 28090 | *Acinetobacter lwoffii* | 89 | 1 | 126 |
| 469 | *Acinetobacter* | 78 | 0.99 | 123 |
| 13690 | *Sphingobium yanoikuyae* | 60 | 0.99 | 135 |
| 645687 | *Astrovirus VA1* | 46 | 1 | 46 |
| 133448 | *Citrobacter youngae* | 37 | 0.91 | 169 |
| 199310 | *Escherichia coli CFT073* | 33 | 1 | 35 |
| 56946 | *Afipia broomeae* | 28 | 1 | 77 |
| 618 | *Serratia odorifera* | 23 | 0.92 | - |
| 409438 | *Escherichia coli SE11* | 21 | 0.98 | 49 |
| 1747 | *Propionibacterium acnes* | 13 | 0.97 | 35 |
| 1282 | *Staphylococcus epidermidis* | - | - | 10 |
| 28211 | *Alphaproteobacteria* | - | - | 10 |
| 28037 | *Streptococcus mitis* | - | - | 8 |
| 562 | *Escherichia coli* | - | - | 8 |
| 509173 | *Acinetobacter baumannii AYE* | - | - | 7 |
| 41297 | *Sphingomonadaceae* | - | - | 6 |
| 40214 | *Acinetobacter johnsonii* | - | - | 6 |
| 29391 | *Gemella morbillorum* | - | - | 5 |
| 76122 | *Alloprevotella tannerae* | - | - | 4 |
| 652103 | *Rhodopseudomonas palustris DX-1* | - | - | 2 |
| 268747 | *Prochlorococcus phage P-SSM4* | - | - | 2 |

The presence of the astrovirus was confirmed using real-time RT–PCR. Genome sequencing of the astro– virus in the sample and subsequent study of the consensus sequence showed that we had in fact identified a novel virus, closely related to the VA1 strain ((Brown et al., 2014), in press).

### Pathoscope

Pathoscope identified 22 taxa, corresponding to 15 species and some genera or families (Table 4). It also assigned all 46 reads to the *Astrovirus VA1*. Almost all the species identified from metaMix were identified by Pathoscope, with an additional 9 taxa supported by few reads. As the method can only properly work with unassembled sequence data, an extra BLASTx similarity step had to be performed for the 91,516 reads that had contributed to the 679 assembled contigs. Pathoscope produced the results in less than one minute.

### MEGAN

MEGAN identified 19 taxa and did not detect the astrovirus signal. We modified the minimum read support parameter from 50 reads to 10 to increase sensitivity. MEGAN then identified 25 taxa, including the *Astrovirus VA1.* The remaining 24 were mostly genera, relevant to the species detected by metaMix and Pathoscope. MEGAN produced the results in less than one minute.

## Human Clinical Sample – Species Absent from the Database

We then compared the performance of the three methods in a scenario where sequences present in the sample are absent from our reference database.

We analyzed a second brain biopsy sample from an undiagnosed patient. 32 million RNA-Seq reads were obtained using the HiSeq instrument. Following initial processing using our bionformatics pipeline, the dataset had 1,261,575 non host reads and contigs for subsequent analyses. There were 3,150 potential species based on the BLASTx output.

### metaMix

The resulting species profile consisted of 7 species (Supplementary Table S2). The most interesting finding was the identification of a human coronavirus. We found two different strains: *Human coronavirus OC43* had almost a million reads assigned to it. Additionally there were 67K reads assigned to *Human enteric coronavirus strain 4408*. The presence of both viral strains in the results indicated that even though the virus in the sample was mostly similar to the OC43 strain, there were sequences sharing higher similarity to 4408 at some loci. This is highlighting how the database choice impacts the results: the RefSeq database we used has only one OC43 strain, while in GenBank there are several OC43 strains capturing the high mutation rates of the *Human Coronavirus OC43*.

We followed up on the sequences assigned to the “unknown category”, that is approximately 170K reads, looking for nucleotide similarity with NR–NT using BLASTn. Approximately half of the reads originated from an untranslated region of the Coronavirus genome, which is not captured by the protein reference database. The remaining reads matched confidently to either *Danio rerio* (zebrafish) sequences or *Gallus gallus* (chicken), two organisms whose proteins are not in the human microbiome reference we are using. The zebrafish and chicken matches were explained as barcode leakage resulting from multiplexing on the same flowcell zebrafish and chicken RNA–Seq libraries. metaMix appropriately assigned these reads to the “unknown” category, producing a clean probabilistic summary (Supplementary Table S2). The method ran in 4.7 hours.

In this instance, the metaMix results emphasize the importance of being able to deal with missing reference sequences that do not have a closely related strain or species in the same database.

### Pathoscope

Pathoscope identified 177 species in this sample (Supplementary Table S2). We optimized the value of the unique read penalty parameter and we achieved the best results with the thetaPrior parameter set within the range 10–100. With these settings, the method identified 52 species (Supplementary Table S2). Our assessment is that Pathoscope is confused by the lack of completeness of databases combined with the absence of an “unknown” category, which prevents it from dealing with these unassigned reads sensibly. Pathoscope completed its analysis in 10 minutes.

### MEGAN

MEGAN assigned the reads to 30 taxa. These included some species and genera but most were families (Supplementary Table S2). Approximately 250K reads could not be assigned to any taxonomic level. MEGAN run in 8 minutes.

## Discussion

Here, we present metaMix, a sensitive method for metagenomic species identification and abundance estimation. The method is implemented in an R package (http://cran.r-project.org/web/packages/metaMix). Using a Bayesian mixture model framework, we account for model uncertainty by performing model averaging and we resolve ambiguous assignments by considering all reads simultaneously. A key feature of the method is that it provides probabilities that answer pertinent biological questions, in particular the posterior probability for the presence of a species in the mixture. Additionally it accurately quantifies the relative proportions of the organisms.

This general framework is designed to address interpretation issues associated with closely related strains in the sample, low abundance organisms and absence of genomes from the reference database. We show that metaMix outperforms other methods in the community profiling task, particularly when complex structures with closely related strains are studied. As a consequence, it also produces more accurate relative abundance estimates for the species in the mixture. The method can deal with either unassembled reads or assembled contigs or both, allowing for flexibility of choice for the bioinformatics preprocessing. In practice, the choice of bioinformatics processing prior to the application of our Bayesian mixture analysis must be optimized for each application, and our processing pipeline has been designed with viral sequence identification from transcriptome sequencing as a main goal. Nevertheless, as demonstrated by our analysis of the mock bacterial community dataset, the method can be applied in other contexts.

The sensitivity and general applicability of metaMix comes at an increased computational cost, requiring access to a multi–core computer to run efficiently. For the datasets presented here, the computation time remained manageable and did not exceed a few hours, using 12 cores to run 12 parallel chains. Nevertheless, a limitation of metaMix is the increased processing time for very large datasets. Speed related improvements can be implemented in scenarios where the species ambiguity concerns only a small proportion of the read set. Reads with certain assignments can be flagged prior to the MCMC exploration of the state-space. Their assignment information can then be carried forward, thereby reducing the size of the similarity matrix used as input by the mixture model. Another area of possible improvement is MCMC convergence determination. The current version of metaMix produces log-likelihood traceplots allowing the user to visually inspect the MCMC convergence, however additional diagnostic criteria can be implemented in future versions.

metaMix is most useful for complex datasets for which the interpretation is challenging. It has been mainly used as a clinical diagnostic tool, helping with the identification of the infecting pathogen while providing an accurate profile of the community in the sample.

## Methods

### Bioinformatics Preprocessing

Prior to running the mixture model for metagenomic profiling, several steps are required to process the short read sequence data (Figure 3). The pipeline uses publicly available bioinformatics tools for each preprocessing step.

**Figure 3:**
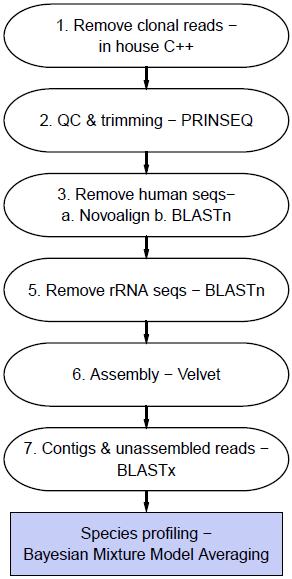
Pipeline steps for species identification. The step of removing the rRNA sequences is applicable only when the aim is viral discovery.

The first step is the removal of clonal reads using an in house C++ script. We then use PRINSEQ (Schmieder and Edwards, 2011) for read-based quality control, removing low quality and complexity reads and performing 3’end trimming. For metagenomic analysis of human samples, reads originating from the human host are not relevant for our research question. We therefore remove human host reads, using a two–step approach to limit computation time: initially a short read aligner (novoalign, www.novocraft.com), followed by BLASTn. The next step is only applicable when the focus is on virus discovery using transcriptome reads. We remove ribosomal RNA sequences, using BLASTn against the Silva rRNA database (http://www.arb-silva.de/).

The remaining reads are assembled into contigs using the Velvet short read assembler (Zerbino and Birney, 2008). For each contig we record the number of reads required for its assembly, using this information at the stage of species abundance estimation. A Velvet tuning parameter is the user defined k-mer length that specifies the extent of overlap required to assemble read pairs. Metagenomic assembly is not a straightforward task, as short k-mers work best with the low abundance organisms, while long k-mers with the highly abundance ones. The shorter the k-mer the greater the chance of spurious overlaps, hence we choose relatively high k-mer length, in order to avoid chimeric contigs.

For each contig and unassembled read we record the potentially originating species, using the nucleotide to protein homology matching tool BLASTx. We use BLASTx due to the higher level of conservation expected at the protein level compared to nucleotides. This choice is guided by our focus in viral pathogens - viruses having high genetic diversity and divergence (Fancello et al., 2012).

This step generates a sparse similarity matrix between the read sequences and the protein sequences, with species as columns, reads and contigs as rows.

The statistical method described in the remainder of this section considers the competing models that could accommodate our observed data, that is the BLASTx results and compares them. The different models represent different sets of species being present in the sample. The method works on two levels of inference: in the first instance we assume a set of species to be present in the sample and we estimate this model’s parameters given the data. The other level of inference is the model comparison so as to assess the more plausible model. The process is iterated in order to explore the model state space.

### Model Specification Assuming a Fixed Set of Species

Assuming a given set of *K* species from which the reads can originate, the metagenomic problem can be summarized as a mixture problem, for which the assignment of the sequencing reads to species is unknown and must be determined. The data consist of *N* sequencing reads ***X*** = (*x_1_,…, x_N_*), and for a given read *x_i_* the likelihood is written as: 
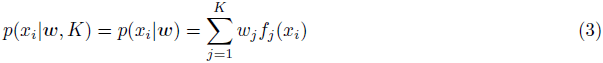
where *w* = (*w_1_, …, w_K_*) represent the proportion of each of the *K* species in the mixture. These mixture weights are constrained such that 0 ≤ *w_j_* ≤ 1 and ∑*_j_ W_j_* = 1. In practice, we also add a category (species *K* + 1) which we refer to as the “unknown” category, and captures the fact that some reads cannot be assigned to any species.

Additionally *f_j_(x_i_) = P(x_i_|x_i_* from species *j*) = *p_ij_* is the probability of observing the read *x_i_* conditional on the assumption that it originated from species *j*. We model this probability using the number of mismatches *m* between the translated read sequence and the reference sequence and a Poisson distribution with parameter λ for that number of mismatches. 
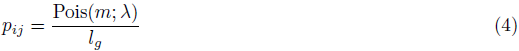
where *l_g_* is the length of the reference genome, when short reads are matched to a nucleotide database. For nucleotide matching, *l_g_* has a large impact on the probability computation. However, when matching against protein databases, the more limited heterogeneity of protein lengths results in a much smaller impact of the length parameter. In addition, incomplete annotation can potentially make the inclusion of protein length problematic for the *p_ij_* computation. Consequently, for protein matched sequences, we simply defined our *p_ij_* as: 
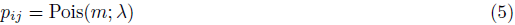

Therefore for a given set of *K* species, the *p_ij_* probabilities are regarded as known and the mixture weights must be estimated. Combining the above we see that when we know the set of species *K*, the mixture distribution gives the probability of observing read 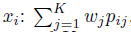, namely equation (3).

We therefore write the likelihood of the dataset ***X*** as a sum of *K^n^* terms: 
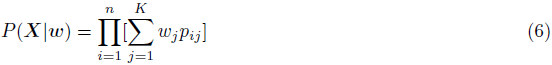

### Estimation of Mixture Weights

Assuming a fixed set of species, the posterior probability distribution of the weights w given the read data *X* is: 
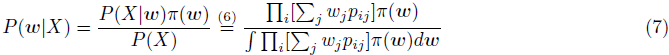

A practical prior for the mixing parameters *w* is the Dirichlet distribution owing to its conjugate status to the multinomial distribution. Despite the use of conjugate priors, the probabilistic assignment of reads to species involves the expansion of the likelihood into *K^n^* terms which is computationally infeasible through direct computation. An efficient estimation can be performed by the introduction of unobserved latent variables that code for the read assignments. In this framework, either the Gibbs sampler (Marin et al., 2005), a Monte Carlo Markov Chain technique, or the Expectation-Maximization (EM) (Dempster and Laird, 1977) algorithm can be used to estimate the mixture weights *w*. EM returns a point estimate for *w* while the Gibbs sampler the distribution of *w* (Supplementary Text 1 for details of implementation). Both methods were implemented and provided comparable results.

### Marginal Likelihood Estimation

Each combination of species corresponds to a finite mixture model for which the marginal likelihood can be estimated. Marginal likelihood comparison has a central role in comparing different models {*M_1_,…, M_m_*}. To compute the marginal likelihood *P* (*X | M_k_*) for the mixture model *M_k_* one has to average over the parameters with respect to the prior distribution *π*(*θ_k_|M_k_*), where *θ_k_* are the model parameters: 
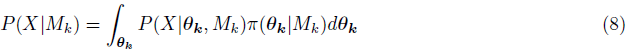

The posterior probability of the model *M_k_* is: 
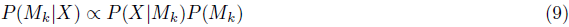
where *P*(*M_k_*) is the prior belief we hold for each model. The prior can be specified depending on the context but the basis of our interpretation is that parsimonious models with a limited number of species are more likely. Thus in this Bayesian framework our default prior uses a penalty limiting the number of species in the model.

We approximate this penalty factor based on a user-defined parameter r that represents the species read support required by the user to believe in the presence of this species. We compute the logarithmic penalty value as the log-likelihood difference between two models: one where all *N* reads belong to the “unknown” category and one where r reads have a perfect match to some unspecified species and the remaining *N* — r reads belong to the “unknown” category. In the nucleotide similarity situation, the *p_ij_* probabilities for the r reads originating from this unspecified species are approximated by 1/(median genome length in the reference database). This parameter essentially reflects how many reads are required to provide credible support that a species is present in the mixture and acts as a probabilistic threshold as opposed to a deterministic one applied on a ranked list.

From now on, when we refer to the marginal likelihood, we mean the marginal likelihood for a specific model and we forego conditioning on the model *M_k_* in the notation. Additionally, in our mixture model *p_ij_* are always regarded as known, therefore the model parameters *θ_k_* are the mixture weights *w*. Hence (8) becomes: 
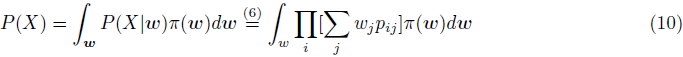

Approximating the marginal likelihood is a task both difficult and time-consuming. We chose the Defensive Importance Sampling technique (Hesterberg, 1995) for the relative simple implementation compared to other approaches (Supplementary Text 1 for details of implementation). This is crucial as we perform this approximation numerous times, for every species combination we consider.

However the goal of this work is to deliver results in a clinical setting within an actionable time-frame. We wish to speed up the computation without compromising the accuracy and the sensitivity of the results. For that reason, we use a point estimate of the marginal likelihood by means of the Expectation-Maximization (EM) algorithm. The different approaches were used on the benchmark dataset. The resulting taxonomic assignment as well as the species relative abundance estimates were similar between them, with the EM approach resulting in a 13-fold speed increase (Supplementary Text 1).

### Model Comparison: Exploring the Set of Present Species

We use a Monte Carlo Markov Chain (MCMC) to explore the set of present species of size 2*^S^* — 1, where *S* is the total number of potential species. In practice we observe that *S* can be greater than 1,000. The MCMC must explore the state-space in a clinically useful timespan. Therefore we reduce the size of the state-space, by decreasing the number of *S* species to the low hundreds. We achieve this by fitting a mixture model with *S* categories, considering all potential species simultaneously. Post fitting, we retain only the species categories that are not empty, that is categories that have at least one read assigned to them.

Let us assume that at step *t*, we deal with a set of species that corresponds to the mixture model *M_k_*. At the next step (*t* + 1), we either add or remove a species and the new set corresponds to the mixture model *M_l_*. The step proposing the model *M_l_* is accepted with probability: 
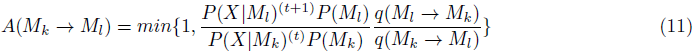
where *q*(*M_l_ → M_k_*) is the probability of transitioning from model *M_l_* to model *M_k_*. In other words, this is the probability of adding or removing the species to the *M_k_* set of species that took us to the *M_l_* set of species.

If the step is accepted, then the chain moves to the new proposed state *M_l_*. Otherwise if not accepted, the chain’s current state becomes the previous state of the chain, i.e the set of species remains unchanged.

metaMix outputs log-likelihood traceplots so that the user can visually inspect the mixing and the convergence of the chain. The default setting is to discard the first 20% of the iterations as burn-in. We concentrate on the rest to study the distribution over the model choices and perform model averaging. We can then summarize appropriately the posterior distribution and answer the important questions of interest. Examples of such questions include: what species have probability *p* or greater being included in the set of present species? what is the probability of having the *n* specific closely related strains in the set of present species? Depending on the biological context, one may ask numerous similar or other case-specific questions.

### Optimized Implementation: Parallel Tempering

We observed that simple MCMC does not efficiently explore the complex model state space, as evidenced by the poor mixing of the chain (Figure 4).

**Figure 4:**
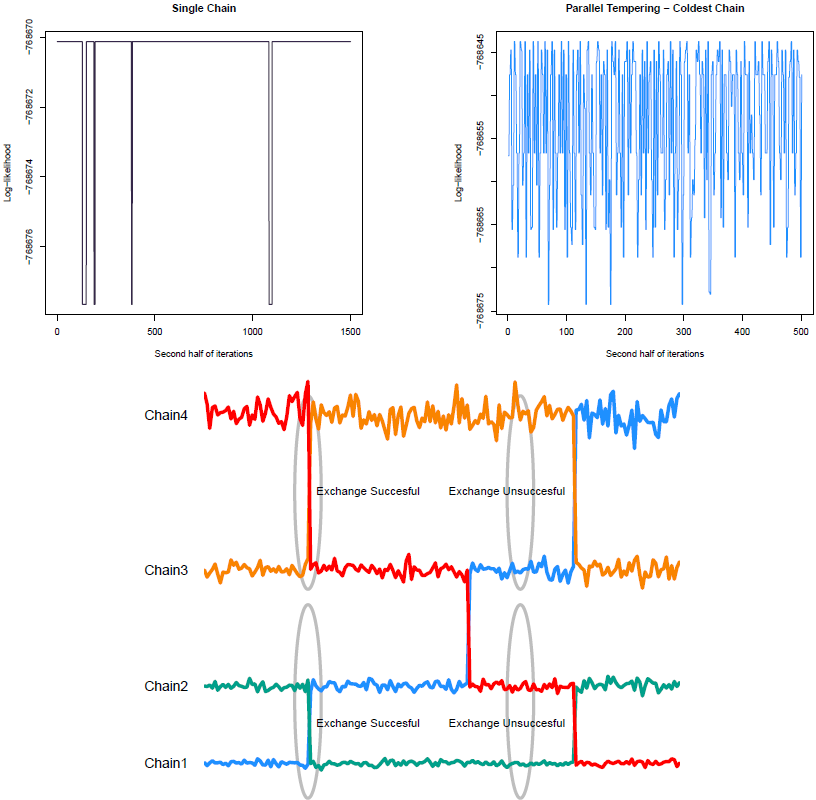
a. Log-likelihood trace plot for single chain MCMC and b. for PT chain at temperature T=1. c. Schematic of parallel tempering. Exchanges are attempted between chains of neighboring temperatures, where Chain1 at *T_1_* = 1, *T_1_ <T_2_ <T_3_ <T_4_*.

In order to overcome this and take advantage of parallel computing, we run multiple chains and allow exchange moves between them. This method is called parallel tempering MCMC (Earl and Deem, 2005). Within the parallel setting, each chain simulates from the posterior distribution *g*(*M*)=*P*(*M|X*) raised to a temperature *t*ϵ(0,1], where model *M* 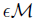 represents a set of species being present. The different temperature levels result in tempered versions of the posterior distribution *P*(*M_k_|X*)*^t=1/T^*. When *T* =1 the draws are from the posterior distribution. On the other hand, at higher temperatures the posterior spreads out its mass and becomes flatter. In practice that means that distributions at higher temperatures are easily sampled, improving the mixing. We are interesting in studying the original posterior distribution with *T* = 1.

We implemented two types of moves. The first is the mutation step, which simply is the within chain move we described in the previous section. This is accepted with probability given by (11). The other is the exchange step, a between chains move. This Metropolis-Hastings move proposes to swap the value of two chains *k* and *k* +1, adjacent in terms of *T*. Suppose that the values of the two chains are *M_k_* and *M*_*k*+1_ respectively, corresponding to two different sets of species. The move is accepted with probability (Jasra et al., 2007): 
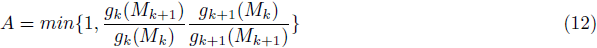

Since *g_k_* (*M_k_*)=*P*(*M_k_|X*)*^T^* and *g*_*k*+1_(*M*_*k*+1_) = *P*(*M*_*k*+1_|*X*)^*T*+1^, it follows that when *M*_*k*+1_ represents a set of species of higher probability than the one *M_k_* represents, the exchange move will always be accepted (Figure 4).

This allows moves between separate modes, ensuring a global exploration of the model state space. Eventually “hot” and “cold” chains will progress towards a global mode.

## Acknowledgments

We thank David Balding for the constructive comments on the manuscript, Christian Robert for the helpful methodological discussion, Judith Breuer, Julianne Lockwood and Mike Hubank for data sharing and informative discussions for the interpretation of the results. SM is supported by a PhD studentship from the Annals of Human Genetics. This work is also supported by the National Institute for Health Research University College London Hospitals Biomedical Research Centre.

